# Novel mechanism of *MYC* deregulation in Multiple Myeloma

**DOI:** 10.1101/2023.05.19.541506

**Authors:** Mahshid Rahmat, Kendell Clement, Jean-Baptiste Alberge, Romanos Sklavenitis-Pistofidis, Rohan Kodgule, Charles P. Fulco, Daniel Heilpern-Mallory, Katarina Nilsson, David Dorfman, Jesse M. Engreitz, Gad Getz, Luca Pinello, Russell Ryan, Irene M. Ghobrial

## Abstract

*MYC* deregulation occurs in 67% of multiple myeloma (MM) cases and associates with progression and worse prognosis in MM. Enhanced *MYC* expression is known to be driven by translocation or amplification events, but it only occurs in 40% of MM patients. Here, we describe a new mechanism of *MYC* regulation, whereby epigenetic regulation of *MYC* by increased accessibility of a cell-type specific enhancer leads to increased *MYC* expression. We found enhancer activity does not associate with enhancer hijacking events. We identified specific binding of c-MAF, IRF4, and SPIB transcription factors to the enhancer can activate *MYC*. In addition, we discovered focal amplification of this specific enhancer in approximately 4% of MM patients. Together, our findings define a new epigenetic mechanism of *MYC* deregulation in MM beyond known translocations or amplifications and point to the importance of non-coding regulatory elements and their associated transcription factor networks as drivers of MM progression.

## INTRODUCTION

Enhanced expression of the *MYC* oncogene is associated with the initiation and maintenance of many human cancers^1, 2^. Multiple myeloma (MM) is a malignancy of clonal plasma cells, in which *MYC* deregulation is a key event in the progression from precursor stages (monoclonal gammopathy of undetermined significance (MGUS) and smoldering multiple myeloma (SMM) to symptomatic MM in 67% of patients^3–5^. *MYC* structural variants (SV) such as translocations, inversions, deletions, amplifications, chromosome 8 monosomies and trisomies, and mutations at the *MYC* partner, *MAX,* are common genomic events that associate with progression of SMM precursor patients to MM and are found in 41% of newly diagnosed MM cases^6–10^. Translocation and amplification of the 8q24.21 *MYC* locus are known SV that mediate *MYC* deregulation at premalignant stages. *MYC* translocations are found in 3-4% of MGUS or SMM and 15-20% of newly diagnosed MM patients. Most translocations result in the juxtaposition of a stretch enhancer or super-enhancer with the *MYC* locus and a substantial increase in its expression^7, 8, 11, 12^. Immunoglobulin (IG)-encoding genes (*IGH, IGK, IGL*) are the most common translocation partners of *MYC*; however, *MYC* translocations also occur with non-IG partners such as *FAM46C, FOXO3,* and *BMP6*. Among the many *MYC* SVs, *MYC-*Ig translocations are associated with poor patient outcomes (IgL in particular)^4, 6, 7, 11^. DNA and RNA sequencing of SMM and MM patients show that *MYC* dysregulation may also occur in the absence of genetic aberrations^6–8, 10, 13^, suggesting that epigenetic pathways contribute to the deregulation of this oncogene in MM progression.

Here we use the CRISPR interference (CRISPRi) system to define the mechanism of *MYC* epigenetic regulation and explore the transcription factor (TF) circuits that mediate *MYC* expression in the context of an intact locus. We also use whole-genome sequencing (WGS) data from MM patients to assess the genetic aberrations at a newly identified enhancer region and define new molecular mechanisms of *MYC* deregulation in MM pathogenesis.

## RESULTS

### Novel *MYC* non-coding regulatory elements in MM

*MYC* transcript levels are higher in MGUS and SMM patients compared to healthy donors and are higher still upon progression to overt MM (Figure 1a)^14^. To investigate the association of *MYC* expression with genetic alterations in MM patients, we used RNA and whole-genome sequencing (WGS) data from the Clinical Outcomes in Multiple Myeloma to Personal Assessment (CoMMpass) dataset analyzed in a previous study^6^. We examined *MYC* expression levels in 678 total MM patients grouped according to *MYC* genetic status: *MYC* loss, *MYC* gain, *MYC* translocations, *MAX* single nucleotide variants (SNVs), and others (MM cases without *MYC* or *MAX* genetic aberrations). 171 cases (25.22%) had *MYC* Ig- or non-Ig translocations and 60 cases (8.84%) had either trisomy 8 (three copies of the *MYC* gene), *MYC* amplifications, or gains (all patients with copy number log2 ratio > 0.1 on the *MYC* locus that did not have *MAX* mutations or *MYC* translocations). Thirty-one cases (4.57%) had monosomy 8 (one copy of *MYC* gene) or *MYC* deletions and 17 cases (2.5%) carried non-silent *MAX* SNV’s which have been hypothesized to functionally substitute for high *MYC* expression in MM pathogenesis ^8, 10, 15^. However, 58.84% of MM patients in the CoMMpass cohort (399 cases) had neither *MYC* SVs or CNAs, nor *MAX* mutations. *MYC* expression was significantly higher in patients with *MYC* rearrangements compared to *MAX* mutated cases, *MYC* loss, or others (*MAX* SNVs versus others, *P* = 1.6E-7; *MYC* loss versus others, *P* = 0.72; *MYC* gain versus others, *P* = 3.6E-7; *MYC* translocations versus others, *P* < 2.22E-16; *MYC* translocations versus *MYC* gain, *P* = 0.013; MAX SNVs versus *MYC* loss, P = 6.3E-7) (Figure 1b). Given the enhanced *MYC* expression in the majority of MM patients and the fact that no *MYC* rearrangement was found in more than 50% of the cohort, we defined MM patients with *MYC* expression greater than the median of the cohort as ‘high *MYC’* patients (log_2_ (1+FPKM) > 5.8; n=339, 50% of participants). Median split is a robust and widely used way to turn a continuous variable into a dichotomous variable. We grouped high *MYC* patients by their *MYC* genetic status; 139 patients (41%) had a translocation of *MYC*, 47 cases (14%) had a *MYC* CNA, and 152 MM patients (45%) had no *MYC* rearrangements (Figure 1b). MM cases with unknown mechanism of *MYC* activation substantially overlapped in their *MYC* expression with cases that had *MYC* rearrangements, suggesting the presence of an uncharacterized mechanism of *MYC* activation. Thus, we hypothesized that epigenetic mechanisms might contribute to *MYC* deregulation in newly diagnosed MM patients with intact *MYC* loci.

**Figure 1.**
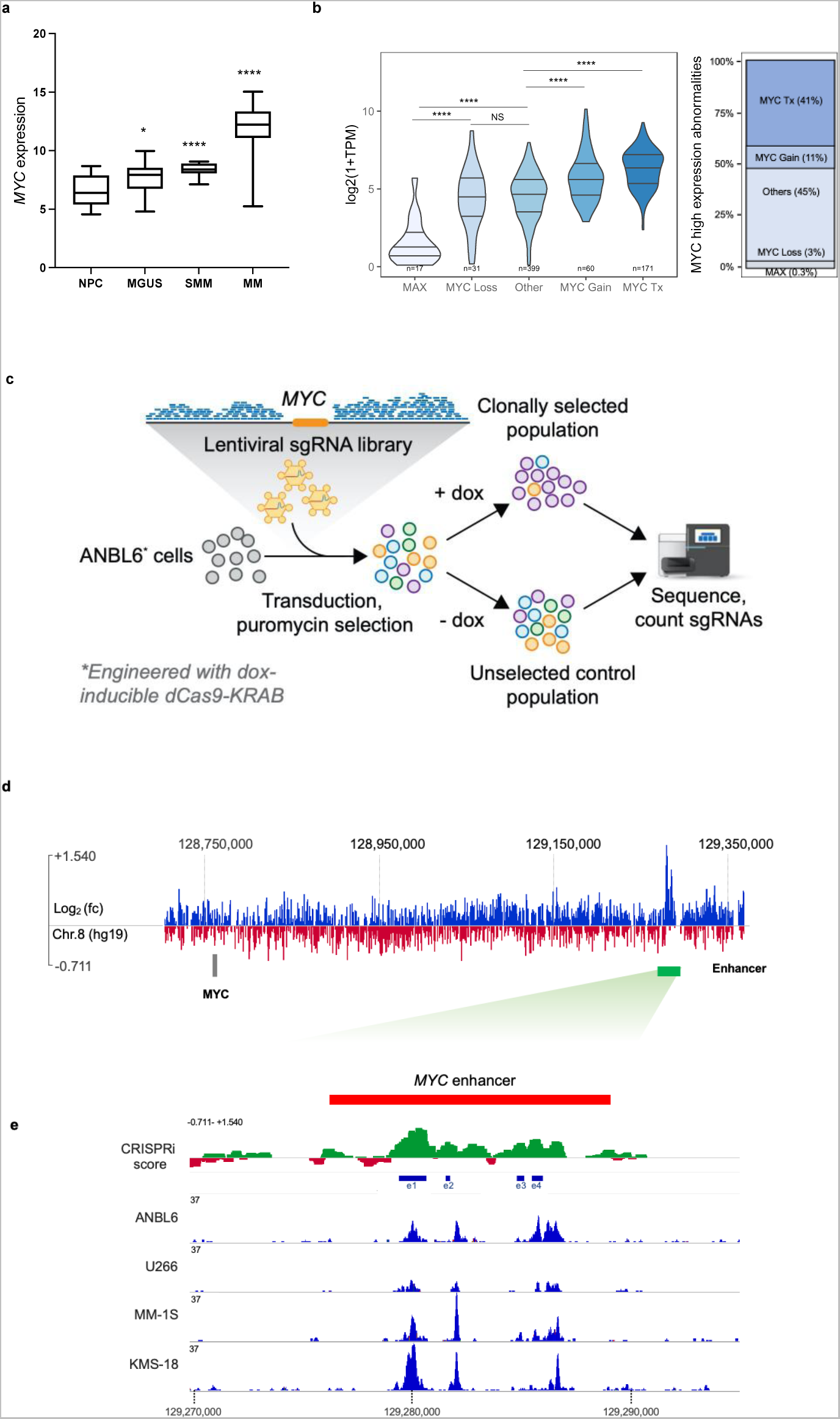
Identification of MYC regulatory elements in MM cells. **a)** MYC expression in Log 2 (1+FPKM) units in normal plasma cells (NPC) from healthy donors, MGUS, SMM, and newly diagnosed MM patients. Each group was compared to NPC with t-test. * P < 0.05, **** P < 0.0001. **b)** (Left) MYC expression in CoMMpass MM patients with MAX mutations (n=17), MYC loss (n=31), MYC gain (n=60), MYC translocations (Tx, n=171), or Other (no known MYC rearrangements, n=399). (Right) MYC high MM patients (MYC expression levels higher than the median; n=339) grouped by MYC loci staus, * P < 0.0001 in t-test, **c)** Schematic representation of the CRISPRi screen. ANBL6 cells expressing KRAB-dCas9 from a doxycycline-inducible promoter were infected with a sgRNA library targeting a total of 1.2 Mb genomic sequence in the MYC TAD as well as 85 kb of control regions. Doxycycline-treated cells were cultured for 14 doubling passages and then subjected to deep sequencing. **d)** Plot of CRISPRi scores for each sgRNA targeting MYC locus and enhancer region in pool screen. **e)** ChIP-sequencing results for H3K27Ac on ANBMl6, U266, MM1S, and KMS-18 MM cell lines. showed the abundance of the histone mark at the enhancer regions.

To identify *cis*-regulatory elements of *MYC* in MM, we conducted a high-throughput CRISPRi tiling screen^16^. We infected ANBL6 cells, which harbor no *MYC* genetic aberrations (Figure 1c), with a lentiviral library of >111,000 sgRNAs tiling across ∼1.2 Mb of DNA around *MYC*. We induced expression of KRAB-dCas9 to epigenetically repress putative regulatory elements. KRAB (Kruppel-associated box domain) is a transcriptional repressor fused to a catalytically dead-Cas9 (dCas9) enzyme and was able to repress the sequence of interest defined by sgRNA. We then sequenced the distribution of sgRNAs in the population before and after 14 passages of growth and defined a “CRISPRi score” as the log_2_ fold-change in sgRNA abundance before and after growth. Because the expression of *MYC* impacts cell growth^17^, sgRNAs that reduce the expression of *MYC* reduce cell division and are less abundant after 14 passages of growth, resulting in strongly negative CRISPRi scores. As expected, sgRNAs targeting the promoter of *MYC* showed significant and consistent reductions across biological replicates (Pearson’s *R* = 0.84, Supplementary Fig. 1a). We also observed a ∼13 kb region distal to the *MYC* gene that, when targeted with sgRNAs, significantly reduced cellular proliferation (Figure 1d). The enhancer region was located ∼523 kb downstream of the *MYC* locus and contained 4 subregions of highly negative scoring sgRNAs (hereafter, e1 to e4).

We inspected the histone acetylation of the identified region by chromatin immune precipitation (ChIP)-sequencing targeting histone H3K27ac in ANBL6, U266, MM-1S and KMS-18 human multiple myeloma cell lines (HMCLs) and found the putative enhancer region was enriched in H3K27ac, which is associated with active enhancers (Figure 1e). In addition, analysis of chromatin structure across B-cell lineage from Blueprint (http://dcc.blueprint-epigenome.eu/#/home) and two other available datasets^18, 19^ revealed that the 13kb enhancer region is accessible in naïve B cells and proliferative plasmablasts indicating that the identified regulatory region is a native plasma-cell specific enhancer of *MYC* (Supplementary Fig. 1b).

Several studies have described *MYC* regulatory elements within the *MYC* topologically associated domain (TAD) in different cancer ^20–23^, so we compared the location of the identified enhancer region in MM cells to known *MYC* regulatory elements and found none of the described *MYC* enhancers overlapped with the MM 13kb regulatory region (Supplementary Fig. 1c). Together, analysis of *MYC* regulatory regions in different cancer types and H3K27ac ChIP-sequencing on MM cells suggest that the newly identified *cis*-regulatory element is an active, MM-specific *MYC* enhancer.

### *MYC* enhancer region is more accessible in MM cells without *MYC* genetic aberrations

*MYC* translocations and gain of the 8q24 locus are secondary genomic events that trigger progression from precursor states to overt MM in 40% of patients^4, 6, 8, 11^. We hypothesized that the enhancers we discovered in ANBL6 cells may lead to *MYC* activation even in the absence of such translocations or other SVs. To examine this, we investigated the chromatin accessibility of the enhancer region in MM patients with and without MYC rearrangements using the assay for transposase-accessible chromatin with high throughput sequencing (ATAC-sequencing). we performed ATAC-sequencing on CD138+ primary MM cells isolated from the bone marrow of 5 newly diagnosed MM cases including two newly diagnosed MM cases with a *MYC* rearrangement and three MM cases without) as well as on normal plasma cells obtained from three healthy donors (Figure 2a). We found that chromatin accessibility at the *MYC* distal-regulatory region was significantly higher in the samples without *MYC* rearrangement than the samples with *MYC* rearrangement or control normal plasma cells (e1, *P* = 0.0279 in *MYC* non-rearranged versus NPC and *P* = 0.0198 in *MYC* non-rearranged versus *MYC* rearranged cases; e3, *P* = 0.015 in *MYC* non-rearranged versus NPC and *P* = 0.0609 in *MYC* non-rearranged versus *MYC* rearranged; e4, *P* = 0.0076 in *MYC* non-rearranged versus NPC and *P* = 0.0716 in *MYC* non-rearranged versus *MYC* rearranged). Of note, chromatin accessibility of e1 and e4 enhancers was comparable in NPC and *MYC*-rearranged patients (e1, *P* = 0.71 in *MYC*- rearranged versus NPC; e4, *P* = 0.706 in *MYC*-rearranged versus NPC) (Figure 2b).

**Figure 2.**
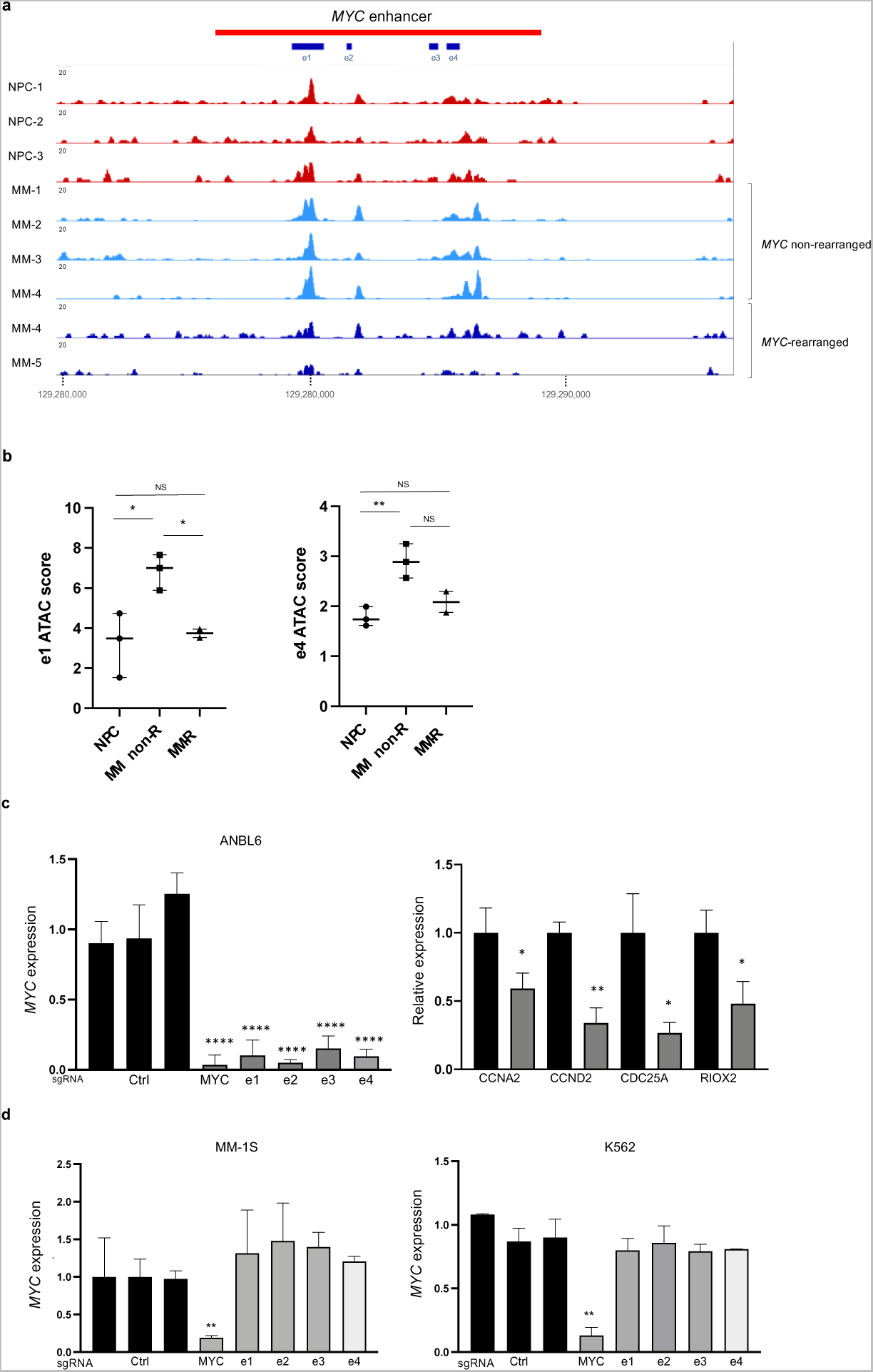
Characterization of *MYC* enhancer elements in MM. **a)** Increased chromatin accessibility at the 13kb regulatory region in MM patients with and without *MYC* genetic aberrations when compared to normal plasma cells. ATAC-sequencing data on malignant plasma cells of three newly diagnosed MM patients without *MYC* translocations and two MM patients with *MYC* rearrangements, **b)** ATAC scores for chromatin peaks at the enhancer regions of MM patients with and without *MYC* rearrangements and plasma cells from healthy donors (NPC). ATAC score was computed using Deeptools’ multiBigwigSummary tool, which reports the number of reads in each region, normalized by sample coverage and region size. The ATAC scores for patients in each group was statistically compared using the standard t-test. Error bars represent 95% CI for the mean values of ATAC scores of the enhancer regions. * *P* < 0.05 in *t*-test versus normal plasma cells. **c)** (left) *MYC* expression by quantitative real-time PCR in ANBL6 cells expressing individual sgRNAs. KRAB-dCas9 expression was induced for 48 hours and values plotted for each sgRNA represent of three independent experiments. Gray bars: sgRNAs targeting one of four enhancers (e1 to e4). MYC: individual sgRNAs targeting the *MYC* TSS (right). Transcript levels (mean with 95% CI) of *MYC* target genes, CCNA2, CCND2, CDC25A and RIOX2, in ANBL6 cells after sgRNA induction (gray bars) were also significantly reduced relative to ANBL6 cells before sgRNA induction (black bars). For each gene, mean value of independent clones plotted carrying sgRNAs against each enhancer element **d)** Suppression of *MYC* enhancer elements in MM1S cells with *MYC* translocation and in K562 non-MM cell lines did not affect *MYC* expression. MYC: sgRNAs targeting the *MYC* TSS. Ctrl: sgRNAs targeting no sequences on the genome. *\* P* < 0.05, *\*\* P* < 0.01, *\*\*\*\* P* < 0.0001 in t-test versus Ctrl sgRNAs for *MYC* expression plots and t-test for *MYC* target gene transcripts in ANBL6 after dox induction of enhancer suppression versus before.

We next targeted each enhancer element with individual sgRNAs in ANBL6 non-*MYC*-rearranged cells. This resulted in an average 89% (*P* < 0.0001) reduction in *MYC* mRNA levels 48 hours after activating KRAB-dCas9 in different sgRNA clones. We also observed significant reductions in the expression of genes targeted by *MYC* as a transcription factor (*CCNA2*, *CCND2*, *CDC25A,* and *RIOX2*^24, 25^) in ANBL6 cells after 48 hours (Figure 2c). These results indicate that the identified non-coding region regulates transcription of *MYC* in MM.

To further assess the functional selectivity of these enhancer elements in the genetically intact *MYC* locus, we measured *MYC* transcript levels in MM1S (*MYC*-rearranged, MM) and K562 (*MYC*-rearranged, non-MM) cells, 48 hours after targeting the e1 to e4 elements. Suppressing the enhancers did not significantly affect *MYC* transcript levels in MM1S and K562 cells as compared to the 89% reduction in ANBL6 non-*MYC* rearranged MM cells (Figure 2d). These findings suggest that the novel enhancer region can activate *MYC* expression in the absence of *MYC* chromosomal translocations in malignant plasma cells.

### IRF4 and MAF are involved in the activation of enhancer elements in non-translocated *MYC*

We next analyzed the sequences of the identified *MYC* enhancers to define binding motifs for transcription factors. Indeed, we identified binding motifs for multiple TFs, including IRF4, cMAF, RELA, POU2F2, MEF2C, SPIB and CTCF. We found one cMAF and one SPIB motifs on e1 and one MAF and two IRF4 binding sites on e4 (Supplementary Fig. 2a). Given the frequency of IRF4 and cMAF and SPIB motifs at the enhancer elements and their known functions in regulation of *MYC* expression, in plasma cell differentiation and MM pathogenesis^26, 27^,we pursued the impact of those TFs in function of newly identified enhancer region. We first assessed IRF4 and cMAF binding to *MYC* enhancer regions by using ChIP-sequencing in *MYC* rearranged cell lines (MM1S and KMS-12) and a non-*MYC* rearranged cell line (ANBL6), (Figure 3a, 3b). Interestingly, there was greater binding of IRF4 to the *MYC* enhancer region in the non-rearranged cell line ANBL6 compared to the rearranged cell lines KMS-12 and MM1S cells, consistent with the functionality of the enhancer region in non-*MYC*-translocated MM cell lines (Figure 3a). MAF bound only at *cis*-regulatory elements of ANBL6 *MYC* non-rearranged cells (Figure 3b).

**Figure 3.**
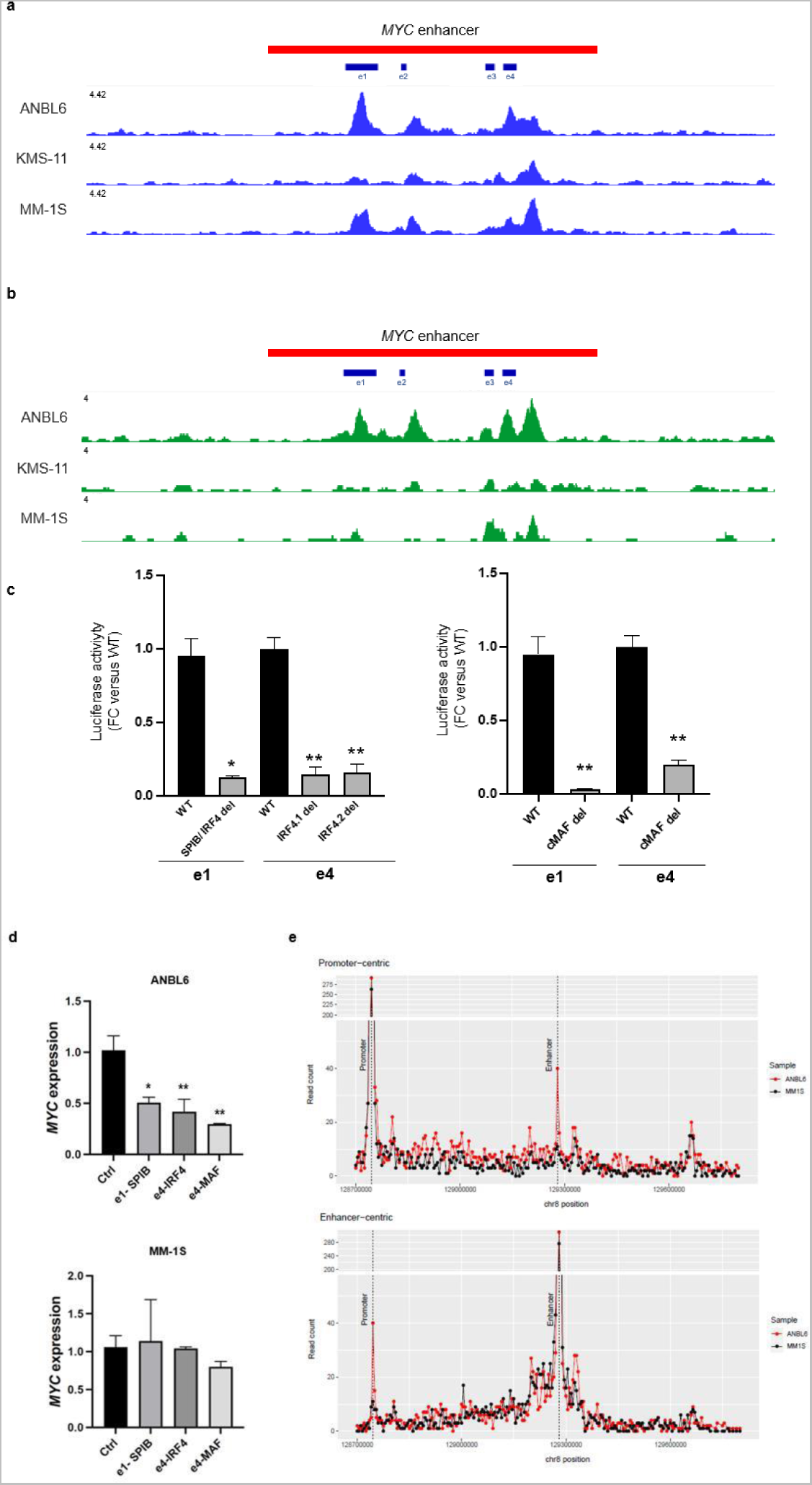
IRF4 and cMAF TFs are involved in *MYC* enhancer activation in MM. ChIP-sequencing for IRF4 **(a)** and cMAF **(b)** in ANBL6, KMS-11, and MM1S cells displayed binding of those TFs to the *MYC cis*-regulatory region with higher abundances in ANBL6 cells that do not carry *MYC* rearrangements in comparison with KMS11 and MM1S translocated cell lines. **c)** e1 and e4 enhancer sequences with and without binding sites for the IRF4 and cMAF were cloned upstream of *MYC* promoter that controls expression of a luciferase reporter gene and luciferase expression signal was measured in ANBL6 cells 24 hours after transfection with WT or deleted reporter constructs. Luciferase signal values were normalized on WT controls and plotted for each construct in biological triplicate. Black bars: WT enhancer constructs, gray bars: enhancer constructs without MAF or IRF4 binding sites. *\*P* < 0.05, *\*\*P* < 0.01, *\*\*\*\*P* < 0.0001 in t-test versus WT controls. **d)** Mutagenesis of TF bindings sites at the enhancer elements of ANBL6 cells. sgRNAs targeting the bindings sites of IRF4, cMAF, and SPIB on e1 and e4 elements were transiently transfected into SpCas9 positive ANBL6 cells and MYC expression was measured 24 hours after. Ctrl: sgRNAs targeting no regions in the genome. *MYC* expression values were normalized on Ctrl sample, *\*\*\*P* < 0.001 in t-test versus controls. **e)** Hi-C chromatin contact frequencies in ANBL6 and MM1S cell lines. The promoter-centric plot (top) shows the number of reads in 5kb bins that support a contact between the *MYC* promoter and genomic loci in the promoter-enhancer TAD in ANBL6 (red) and MM1S (black) cell lines. The lower plot shows the number of reads that support a contact between the *MYC* enhancer and genomic loci in the promoter-enhancer TAD. The genomic positions of the promoter and enhancer are marked with a vertical dotted line.

We next asked whether these transcription factors function as activators. We used a luciferase reporter with wild type (WT) e1 and e4 enhancer sequences or enhancers with the MAF or IRF4 binding sites deleted. The enhancers were upstream of a *MYC* promoter driving luciferase expression. Comparing the expression level of luciferase regulated by WT or mutant enhancer sequences revealed that deletion of the MAF binding site in e1 led to almost no luciferase expression (97% reduction, *P* = 0.0083), and deletion from e4 led to a 79% reduction (*P* = 0.0052). Luciferase expression was also significantly reduced in e1 and e4 enhancers without binding sites for IRF4 (an average of 88% in e1 and 86% in e4, *P* = 0.0102 and 0.0057, respectively (Figure 3c).

Depletion of IRF4 and cMAF in ANBL6 cells led to almost complete (96%) and 70% loss of *MYC* expression (Supplementary Fig. 2b). To examine the contribution of the identified TFs in enhancer activity endogenously, we targeted IRF4, c-MAF, and SPIB binding sites at the e1 and e4 enhancer elements of ANBL6 cells using CRISPR knock-out approach. SPIB is a lymphocyte lineage-specific ETS TF, that is essential for the survival of mature B cells and represses plasma cell differentiation^28–30^. SPIB is also a co-factor of IRF4 in diffuse large B cell lymphoma^31^ and has been described to act as both tumor suppressor and oncogene by activating distinct signaling pathways such as NFkB, JNK, and TGF-beta pathways^32, 33^.

IRF4 and cMAF are important TFs in MM biology and targeting those TFs affect cell survival and proliferation^4, 27, 34^. To assess the role of cMAF, IRF4 and its co-factor SPIB on *MYC* enhancer activity, we targeted each TF binding site at e1 and e4 regulatory elements with individual sgRNAs using a CRISPR KO approach. ANBL6 *MYC* non-rearranged and MM1S *MYC*-rearranged Sp-Cas9 positive cells were transiently transfected with individual sgRNAs and *MYC* expression was measured after 48 hours. Analysis of expression values in pool of targeted cells revealed a 58% decrease (55% IRF4, *P* = 0.0072; 53% SPIB, *P* = 0.016; 70% cMAF, *P*= 0.0019) in *MYC* mRNA levels in ANBL6 cells, while mutated MM1S cells showed comparable *MYC* transcript levels to the control sgRNA cells. These results were in concordance with our luciferase reporter assay, where we showed that deletion of MAF and IRF4 binding sites from e1 and e4 enhancer elements led to almost complete loss of luciferase expression in ANBL6 cells with active *MYC* enhancer (Fig. 3c). Our findings indicated that not only targeting each single enhancer element within 13kb regulatory region, but also mutating single binding sites of enhancer-associated TFs of cMAF, IRF4 and SPIB could interfere with activation of enhancer and induction of *MYC* expression in non-rearranged MM cells.

To examine the interaction of the identified *MYC* enhancer region with the promoter, we profiled genome topology by Hi-C on MM1S and ANBL6 cells. We used HiCCUPS^35^ to identify chromatin contacts genome-wide. Overall, 12,979 and 11,116 contacts were found in the ANBL6 and MM1S samples, respectively. We examined the three-dimensional architecture surrounding the *MYC* promoter and observed several topological interaction domains starting at the *MYC* promoter and extending up to 2Mb downstream from the *MYC* promoter, including the identified *MYC* enhancer region (Supplementary Fig. 3). Within the *MYC* promoter-enhancer domain, HiCCUPS identified six significant contacts in ANBL6 (including the *MYC* promoter-enhancer contact) and two significant contacts in MM1S. In ANBL6, the *MYC* promoter-enhancer contact was supported by 40 reads (at 5kb resolution), and significant in all comparisons at a corrected *p*-value of at least 2.76e-15. (Fig. 3e and Supplementary Fig. 3). We counted the number of reads in 5kb tiles in the *MYC* promoter-enhancer domain and found that the *MYC* promoter interacted most frequently with the identified *MYC* enhancer region (after excluding *cis* interactions within 25kb) in ANBL6 (Fig. 3e and Supplementary Fig. 3).

Collectively, our findings indicate that *MYC* deregulation in malignant MM plasma cells can be regulated by a third mechanism, whereby, the selective gain of chromatin access at the enhancer region augments the transcription of *MYC* via recruitment of IRF4 and cMAF regulatory factors and bring the identified enhancer region in contact with the *MYC* promoter in the absence of *MYC* chromosomal abnormalities.

### Focal amplification of the enhancer elements in MM patients

Given that the *MYC* enhancer is located within the genomic region that is rearranged in MM patients, we asked whether the enhancer region can also be amplified in myeloma. We examined the translocation breakpoints and duplications that occurred within the *MYC* TAD domain of 892 newly diagnosed MM patients using long-insert whole-genome sequencing (WGS) data from the CoMMpass study. *MYC-IG* and non-*IG* translocations were found in 203 MM patients (22.76%), tailing the *MYC* locus, including the enhancer region. This is in concordance with previous studies of CoMMpass data^7, 8^. However, a heatmap of copy number alterations across the 2.9 Mb *MYC* TAD revealed that duplications within 13kb of the identified enhancer region occurred in 3.81% of the MM patient cohort (n=34 patients). In 88.2% of the enhancer duplicated patients (n=30), we did not find any other MYC rearrangements at the TAD. Of note, focal amplification of the *MYC* enhancer region was observed exclusively in MM patients that did not have other *MYC* translocations or duplications within *MYC* TAD (Figure 4a).

**Figure 4.**
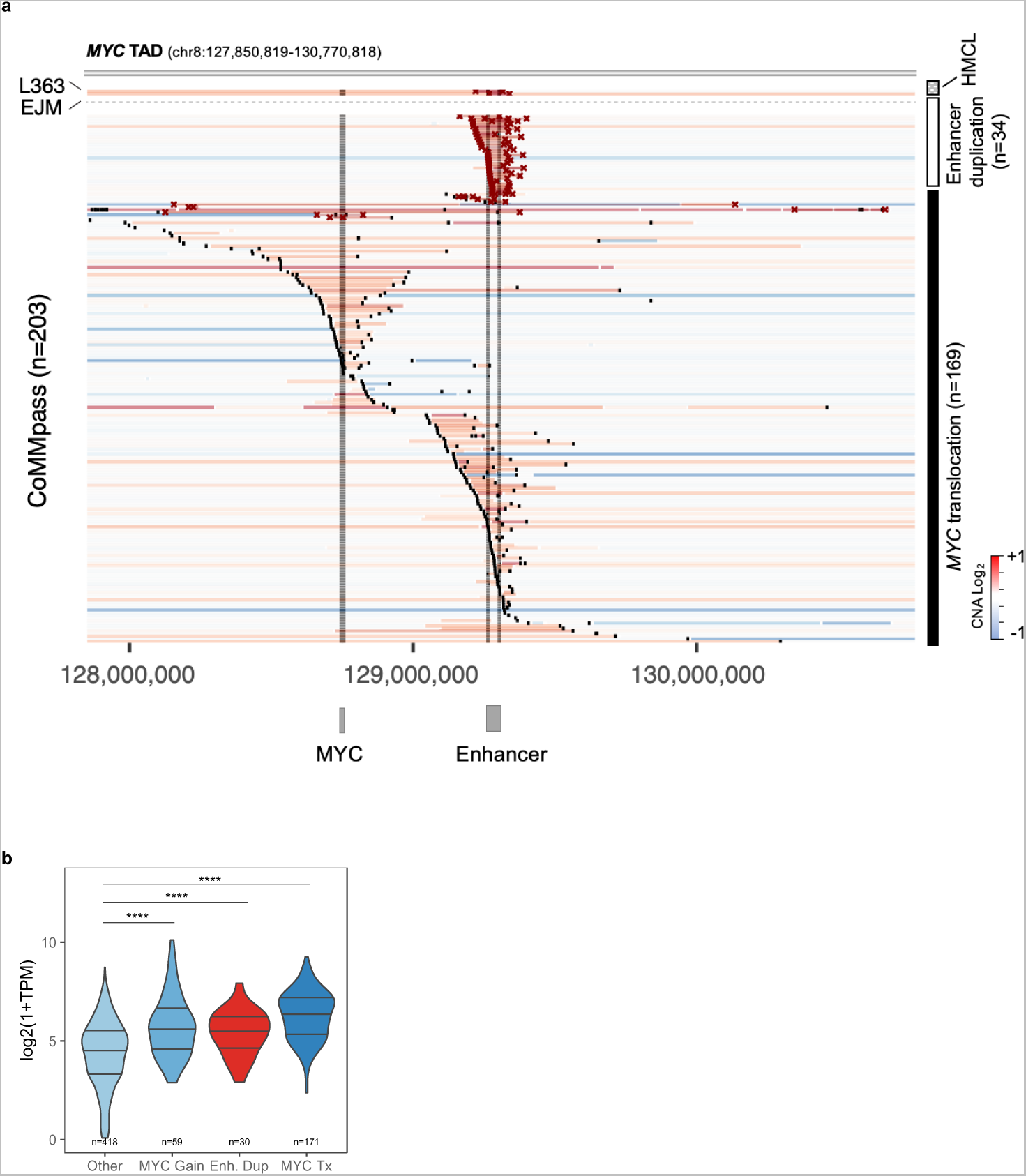
*MYC* enhancer is focally amplified in MM patients. **a)** *MYC* duplications are exclusively found at the *MYC* enhancer sequence. A heatmap of *MYC* CNA across the 2.9 Mb *MYC* TAD for 203 MM patients with *MYC* translocations or enhancer duplication. Each horizontal line represents one MM cell line (EJM or L363) or MM patients with a *MYC* translocation or duplication. Translocation breakpoints and enhancer duplication sequences were plotted as 10kb black and red boxes, respectively. Gray bars display the location of the enhancer region identified in CRISPRi screen and a 25 kb flanking region in enhancer duplicated MM patients and L363 and EJM MM cell lines, **b)** *MYC* expression levels in MM patients without *MYC*–rearrangements (Other), with *MYC* translocations (*MYC*-Tx), MYC gain and patients showing enhancer duplication (Enh. Dup). Focal amplification of the enhancer region led to similar *MYC* transcript levels as translocated patients and a significantly higher *MYC* expression compared to MM patients without *MYC* rearrangements. *\*\*\*\* P* < 0.0001 in t-test versus none.

We then investigated the occurrence of *MYC* enhancer amplifications across 996 different human cancer cell lines from the DepMap portal and identified enhancer gains in only 2 MM cell lines— EJM and L363, further confirming the specificity of the identified enhancer in MM (Figure 4a).

We next investigated the impact of enhancer duplication on *MYC* expression using matched RNA-sequencing data from CoMMpass. *MYC* translocations, gains, and enhancer duplications were all associated with increased *MYC* transcript levels compared to *MYC* non-rearranged cases (*MYC* gains versus others, *P* = 1.4E-8; enhancer duplications versus others, *P* = 2.1E-5; *MYC* translocations versus others, *P* < 2.2E-16) (Figure 4b). Focal amplification of the identified enhancer region is the only duplication event we identified within the *MYC* TAD domain. Amplification of enhancer is a new genetic aberration that did not co-occur with other *MYC* rearrangements but resulted in similar *MYC* expression levels in MM patients.

Together, our findings suggest novel epigenetic and genetic mechanisms of *MYC* deregulation. Selective activation of a newly defined enhancer region either through gain of chromatin access or by genetic amplification induces the expression of this oncogene in the absence of known *MYC* rearrangements in MM patients (Figure 5). We identified a novel genetic mechanism of *MYC* deregulation in newly diagnosed MM patients; in 4% of these cases, *MYC* deregulation stems from duplications found exclusively at the *MYC* enhancer sequence, beyond previously known *MYC* translocation and amplification events.

**Figure 5.**
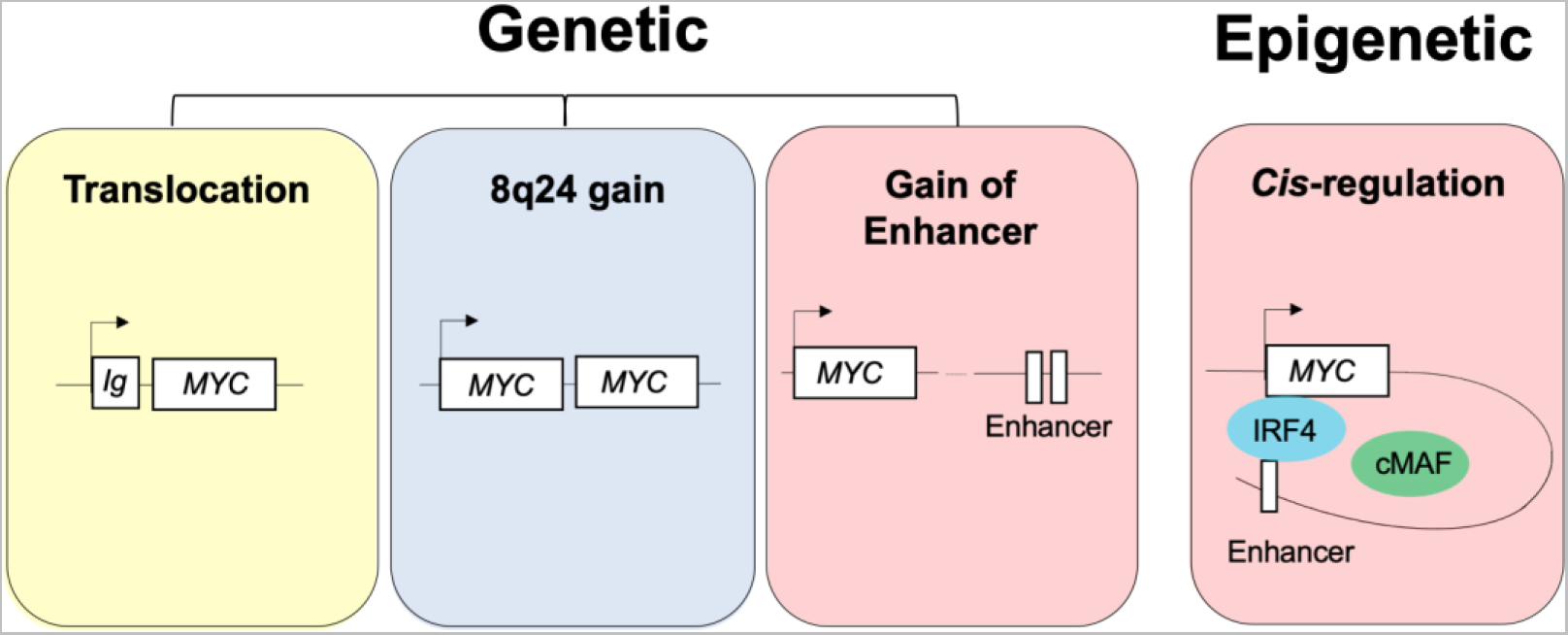
Mechanisms of *MYC* deregulation in MM. Ig- and non-Ig translocations and gain of *MYC* loci (trisomy 8, duplications and amplifications) are known genetic aberrations that lead to enhanced *MYC* expression in MM progression. We have defined an epigenetic mechanism of *MYC* deregulation in which gain of chromatin accessibility at a distal *cis*-regulatory region results in recruitment of MM-specific IRF4 and cMAF TFs and activation of transcriptional machinery to induce the expression of *MYC* oncogene in MM cells. Moreover, we discovered focal amplification of the enhancer region as a novel genetic alteration that leads to the deregulation of *MYC* in 4.13% of MM patients.

## DISCUSSION

Increased *MYC* expression occurs in most MM cases and is associated with progression from precursor stages to overt MM^4, 8^. Genetic studies have described *MYC* structural variations in ∼ 40% of newly diagnosed myeloma patients, with the majority of these consisting of “enhancer hijacking” rearrangements that juxtapose the *MYC* gene to powerful enhancers from distant genomic loci that do not regulate *MYC* in normal development. These rearrangements involve diverse enhancer-rich partner loci, and frequently involve additional co-amplification of both the MYC gene and heterologous enhancer^9–11, 36^.

Here, we used the CRISPRi system to identify an alternative mechanism in which an enhancer native to the *MYC* locus controls *MYC* expression in malignant plasma cells. Inhibiting the *MYC* enhancer elements by CRISPRi led to ∼ 90% reduction in *MYC* transcript levels in non*-*rearranged cells compared to that in *MYC*-rearranged MM cells. The MM enhancer region does not overlap with known *MYC* regulatory regions identified in other cancers, but is active in normal plasmablasts, implying that the subset of myelomas dependent on this native enhancer rely on hyperactivation of the native cell state-specific *MYC* distal regulatory machinery. Genetic events within the MYC locus appear to contribute to this hyperactivation in some cases, as focal amplification of the identified enhancer sequence was the only structural variant in the 2.9 Mb *MYC* TAD domain in 4% of MM patients. Gain of enhancer region led to higher *MYC* expression in the affected individuals compared to non-rearranged patients, but at similar levels to other *MYC* genetic aberrations.

Altered chromatin state of enhancer region provides accessible binding site for cMAF, IRF4, and its co-activator SPIB, well-known transcription factors in MM pathogenesis. IRF4 is a master regulator of B cell development and plasma cell differentiation and an essential gene in MM^26, 27^. Although it is expressed in normal plasma cells, its transcription level steadily increases throughout MM progression^27, 34, 37, 38^. Several studies in B cell malignancies, including MM, describe the binding of IRF4 to the *MYC* promoter and the creation of an autoregulatory feedback loop deregulating these two genes^39–41^. However, this is the first description of how binding of cMAF and IRF4 to a small non-coding regulatory region, changed the chromatin conformation and led to *MYC* activation in non-translocated cell lines.

cMAF is a member of the MAF family of transcription factors and one of the primary translocation events, t(14;16), that is present in about 5-10% of newly diagnosed MM patients^4, 42^. Translocation of c-MAF is associated with unfavorable overall survival in MM^43, 44^. Our data indicate that c-MAF can play an unfavorable role even in non-t(14:16) translocated cells by enhancing the expression of *MYC* in non-*MYC* translocated cells. It also opens the door for potential mechanisms of regulation of *MYC* expression by targeting cMAF or IRF4 in non-translocated cells.

As technical and analytic methods for exome sequencing have become routine, and recurrent driver coding mutations have been exhaustively discovered in common and rare cancer types, one of the most significant remaining challenges in cancer genetics is that of the “missing” drivers - oncogenic lesions that cannot be discovered by such routine methods. Driver lesions in the non-coding genome are still challenging to identify, and descriptions of new discovery approaches and new lesions continue to have a substantial impact^45^. Here, we used the combination of a high-throughput functional CRISPRi screen and high-resolution copy number and WGS data to identify a novel common driver in a common cancer and we expect that it will influence future investigations into other malignancies. We uncovered new epigenetic and genetic mechanisms of *MYC* deregulation whereby, activation of a *cis*-regulatory region through gain of chromatin access and or amplification of the enhancer sequence, provide a platform for binding of cMAF and IRF4 and recruitment of transcriptional machinery to the *MYC* promoter in MM cells. Our results point to the importance of chromatin state in predicting functional enhancers, the impact of genetic alterations at the non-coding regulatory elements on gene expression, and eventually the role of the TF regulatory networks in the deregulation of transcription profiles in human diseases. These novel insights could lead to the identification of new predictive biomarkers and therapeutic targets to benefit patient outcomes in MM and other cancers.

## METHODS

### Cell lines and patient samples

U266, ANBL-6, KMS-12BM, MM-1S, and K562 cells were maintained with RPMI (Thermo Fisher Scientific, Cat # MT10040CV) supplemented with 10% fetal bovine serum (Thermo Fisher Scientific, Cat # 26140079) and 1% pen-strep (Corning, Cat # 30-001-CI). ANBL-6 cells were also supplemented with 5 ng/ml IL-6 (Thermo Fisher Scientific, Cat# 206IL010).

Bone marrow mononuclear cells were collected from SMM and newly diagnosed MM patients as well as healthy donors at Dana-Farber Cancer Institute. The study was approved by the Dana-Farber Cancer Institute Review Board (#14-174) and informed consent was obtained in accordance with the Declaration of Helsinki. CD138+ cells were isolated from bone marrow aspirate samples using Human CD138 MicroBeads (Miltenyi Biotec, Cat # 130-051-301). Selected cells were either viably cryopreserved in 10% dimethyl sulfoxide (DMSO) or used immediately for ATAC-sequencing.

### Establishment of CRISPRi lines

Inducible CRISPRi cell lines were generated by transducing MM cells with lentiviruses carrying TRE-KRAB-dCas9-IRES-BFP and a construct expressing rTA linked by IRES to a neomycin resistance gene for induction of KRAB-dCas9 promoter, as described previously^16^. Transduced cells were selected with 400 µg/ml geneticin (Thermo Fisher Scientific, Cat # 10131035); selected cells were treated for 48 hours with 500 ng/ml doxycycline. Blue fluorescent protein (BFP^+^) cells were selected using fluorescence-activated cell sorting (FACS).

### Pooled CRISPRi screen

Lentivirus production for the *MYC* locus tailing pool was produced as described by Fulco *et al*^16^. Inducible KRAB-dCas9 ANBL6 cells were transduced at the coverage of 1000 transfected cells per sgRNA. Thirty-six hours after infection, transduced cells were selected with 2 µg/ml puromycin for 72 hours. One hundred and fifty million cells were collected as the reference control, and another 150 million cells were treated with 500 ng/ml doxycycline and 2 µg/ml puromycin for 14 population doublings. Genomic DNA was extracted from both reference control and doxycycline-induced cells, using the QIAamp DNA Blood Maxi kit (Qiagen, Cat # 51192). Genomic DNA was amplified for Illumina sequencing using sgRNA library primers as described previously^16^ and was sequenced on the HiSeq2500 platform with 31 bp paired-end reads. Sequencing reads were aligned using Bowtie and the CRISPRi score was calculated as the −log_2_ depletion between the reference control and passage 14 cells for two biological replicates. The mean value of the two replicates was used as the CRISPRi score for each sgRNA.

### CRISPR knock-out cell line

To establish a CRISPR knock-out cell line, ANBL6 cells were infected with lentiviruses carrying pCMV-T7-SpCas9-P2A-EGFP (Addgene, plasmid # 139987) construct and GFP^+^ cells were selected using FACS.

### Cloning individual sgRNAs

For each enhancer element (e1, e2, e3, and e4) scored in the screen, we selected two sgRNAs with high specificity from the screening pool. For TF binding site mutagenesis, sgRNAs were designed using CRISPOR (http://crispor.tefor.net/) software, and guides with high specificity and low off-target scores were selected. Individual sgRNAs were cloned into SgOpti (Addgene, plasmid # 85681) as previously described^16^.

### Single sgRNA knockdown

Single guide RNAs were cloned into SgOpti plasmid and 2 × 10^5^ cells expressing each sgRNA were plated in a 24 well-plate and transduced with 500 ng/ml doxycycline. After 48 hours, cells were harvested in Buffer RLT (Qiagen), and gene expression was measured by quantitative PCR. sgRNA sequences are listed in Supplementary Table 2.

### ATAC-sequencing

50,000-100,000 viable cells were harvested and washed with cold phosphate-buffered saline (PBS). Cells were resuspended in 50 µl of cold lysis buffer (10mM Tris-HCl pH 7.5, 10 mM NaCl2, 3 mM MgCl2, 0.01% NP40) and pelleted. The cell pellet was resuspended in Illumina TD buffer and 2.5 µl of Tn5 enzyme was added to the transposition reaction (Illumina, Cat # FC-121-1030). Nuclei were incubated at 37°C for 60 minutes and fragmented DNA was purified using the MinElute kit (Qiagen, Cat #28206). Library preparation was performed on purified fragments using Nextera DNA Flex Library Prep Kit (Cat #20018704) and libraries were sequenced on the HiSeq2500 platform with 100 bp paired-end reads.

Deeptools^46^ was used to calculate enrichments in each region based on bigwig enrichment tracks for each sample produced using the ENCODE ATAC-seq data processing pipeline (https://github.com/ENCODE-DCC/atac-seq-pipeline)^47^. To compensate for differences in sequencing depth and sample quality, the top 10,000 most significant peaks were called by MACS2^48^, and were selected based on their q-value. These peaks were aggregated for all patient samples and the union of all peaks was derived, such that overlapping peaks between patients would be merged into one larger peak. A binary matrix was created (union peaks by patients) with a value of 1 where the patient peakset had a peak overlapping with that union peak and 0 otherwise. Each sample group (Normal, MM) was compared against the others in a pairwise comparison. For each peak in the union peakset, if all (or all but one) samples from one group had a peak overlapping the union peak, and no (or one) samples from the other group had a peak overlapping the union peak, the peak was included in the most-variable-regions set. To define the TF binding sites at the open chromatin regions of patients, we downloaded GEM peaks from Gene Transcription Regulation Database v19.10 http://gtrd.biouml.org/. These include the transcription factor binding sites inferred from 13,515 ChIP seq of 1,339 transcription factors. We also used the V3 Encode Regulation clustered TF binding peaks, including ENCODE data uniformly processed by the ENCODE Analysis Working Group (https://hgdownloadtest.gi.ucsc.edu/goldenPath/hg19/encodeDCC/wgEncodeRegTfbsClustered/rele aseLatest/wgEnco deRegTfbsClusteredV3.bed.gz)^47^. We overlapped these peaks with the 10K union peakset to identify transcription factors in differentially accessible regions.

### Chromatin Immunoprecipitation

2 × 10^7^ cells for H3K27ac (Active Motif, Cat # 39133) and 3 × 10^7^ cells for IRF4 (Santa Cruz Biotechnology, Cat # SC-48338) and cMAF (Bethyl Laboratories, Cat # A700-045) were harvested and crosslinked with 1% formaldehyde using the TruChIP chromatin shearing kit as per the manufacturer’s instructions (Covaris, Cat # 520154). Chromatin was sheared using Covaris M220 Focused-ultrasonicator resulting in ∼ 200 bp fragments and diluted 1:1 with Covaris IP dilution buffer. The sheared chromatin was incubated overnight with an appropriate antibody or with an Ig isotype control (Santa Cruz Biotechnology, Cat # SC-2343) on a rotator, at 4°C. Antibody-bound chromatin were incubated with protein G Dynabeads for 4 hours rotating at 4°C. Chromatin captured beads were washed five times with RIPA wash buffer (0.1% deoxycholate, 0.1% SDS, 1% Triton x- 100, 10 mM Tris-HCl pH 8.0, 1mM EDTA, 140 mM NaCl), and twice with high salt RIPA wash buffer (0.1% deoxycholate, 0.1% SDS, 1% Triton x-100, 10 mM Tris-HCl pH 8.0, 1mM EDTA, 360 mM NaCl), twice with LiCl wash buffer (250 mM LiCl, 0.5% NP40, 0.5% deoxycholate, 1mM EDTA, 10 mM Tris-HCl pH 8.0) followed by two washes with TE. Beads were resuspended in low SDS elution buffer (10 mM Tris-HCl pH 8.0, 1 mM EDTA, 300 mM NaCl, 0.1% SDS). Eluted DNA was incubated at 65°C for 6 hours in a thermocycler for reverse-crosslinking, and 30 minutes with 1µl RNase A (Thermo Fisher Scientific, Cat # 12091) at 37°C. 2.5 µl Proteinase K (Thermo Fisher Scientific, Cat # AM2546) was added to de-crosslinked DNA and incubated at 37°C for 2 hours. Beads were removed by a magnet and SPRI clean-up was performed on ChIP-DNA. Eluted DNA was used to prepare the sequencing library, using the KAPA HyperPrep kit per the manufacturer’s instructions. (Roche Diagnostics, Cat#: KK8504). Libraries were sequenced on HiSeq2500 with paired-end reads. H3K27ac data were processed according to the chip-seq-pipeline2, and peaks were called using MACS2. The peaks from all cell lines (ANBL6, KMS-18, MM1-S, and U266) were merged and overlapped with differentially accessible regions to identify differentially accessible enhancers.

### qRT-PCR

RNA was isolated using the RNeasy Mini Kit (Qiagen, Cat# 74104) and quantified by Nanodrop (Thermo Fisher). First-strand cDNAs were synthesized using SuperScript™ III (Thermo Fisher, Cat# 18080051). For quantitative PCR, 5-10 ng of cDNA were mixed with Power SYBR Green MasterMix (ThermoFisher, Cat#4367659). Each Ct was measured using the QuantStudio™ 7 Flex Real-Time PCR System. Mean dCt for three technical replicates were calculated and relative RNA levels were estimated and compared to GAPDH control. Primers are listed in Table S1.

### Micro-C

ANBL6 and MM1S cells were harvested and washed once with 1X PBS. Cells were collected by spin down at 300g for 5 minutes and cell pellets were frozen aliquots of 1×10^6^ cells at −80 C overnight. Cells were thawed the day after and resuspended in 1 ml 1X PBS and 10 ul 0.3M DSG (Thermo Fisher Scientific, Cat # A35392) and rotated at RT for 10 minutes. 27ul of 37% formaldehyde was added to the tubes and rotated for 10 minutes at RT. Cells were then collected at 300g for 5 minutes and washed twice with 200 ul of 1X Wash buffer from the Micro-C kit (Dovetail, Cat # 21006). After removing the second wash, 500,000 cells were resuspended in 50ul of 1X Nuclease Digest Buffer and 1.5 ul of MNase enzyme mix and incubated at 22°C for 15 minutes in an agitating thermal cycler set at 1250 rpm. Reactions were stopped by adding 0.5 EGTA and 3ul of 20% SDS and incubated again at 22°C for 5 minutes, 1250 rpm. 1 µg of cell lysates were then subjected to proximity ligation and library preparation following Micro-C protocol. Final libraries for biological replicates of each cell line were sequenced on NovaSeq 6000 S1 with 150 bp paired-end reads. ANBL6 library had 792,341,057 total reads, and after deduplication 314,282,616 reads mapped in cis and 69,220,830 mapped in trans. The MM1S library had 973,642,396 total reads and after deduplication 334,057,407 mapped in *cis* and 95,978,342 mapped in *trans*. Reads were analyzed using the recommended analysis steps (https://micro-c.readthedocs.io/en/latest/). Specifically, reads were aligned to the hg19 genome using bwa version 0.7.17 (https://academic.oup.com/bioinformatics/article/25/14/1754/225615). Ligation junctions were identified using pairtools version 0.3.0 (https://link.springer.com/protocol/10.1007/978-1-0716-1390-0_7) Hi-C contact maps were generated using Juicer Tools version 1.22.01^49^. Chromatin loops were identified using the Juicer tools HiCCUPS and HiCCUPS Diff. Counts of reads supporting chromatin loops around the *MYC* promoter and enhancer were calculated at 5kb resolution, and plotted using R.

### Luciferase reporter assay for *MYC* enhancer activity

To examine the function of identified enhancer elements, we synthesized e1 and e4 DNA sequences (GENEWIZ, Cambridge, MA), either containing WT or deleted TF binding sites. Those sequences were then cloned into the pGL4.23-MYC plasmid (Addgene plasmid # 86461) carrying *MYC* promoter upstream of the luciferase gene. For each construct, we transfected 500,000 ANBL6 cells with 1 ug of reporter plasmid plus 250 ng of Rinella plasmids pRL-SV40 and pGL3 control (Promega) in biological triplicates. Cells were collected 24 hours after transfection, washed with 1x PBS, and lysed in 40 ul Passive Lysis Buffer (Promega). Cell lysates were then subjected to Dual-Luciferase Reporter assay (Promega) following the manufacturer’s protocol. Firefly luciferase activity was normalized to Renilla luciferase activity values and plotted for each construct.

### Data availability

ATAC-seq, ChIP-seq, and Micro-C data were generated as part of this study and data have been deposited with links to BioProject accession number PRJNA791908 in the NCBI BioProject database (https://www.ncbi.nlm.nih.gov/bioproject/). K562 ATAC-seq data from a published study^50^ were downloaded from NCBI (Accession number GSE99173).

## Acknowledgments

The authors would like to acknowledge Anna V. Justis, PhD, a medical writer funded by Dana-Farber Cancer Institute, for editing. This work was partially funded by the Prevention Project Perelman Family Foundation Early Disease Translational Research Program, The Pussycat foundation, and the Dr. Miriam and Sheldon G. Adelson Medical Research Foundation (AMRF).

## Author contributions

M.R., K.C., L.P., and I.M.G designed and M.R. performed the experiments with help from R.K. and S.T.; K.C. performed bioinformatic analyses; C.F.P. and J.E. analyzed the CRISPR data; patient samples and clinical information were collected with help of M.B., D.H., A.P.G., and D.D.; L.P., R.S.P., J.B.A., M.P.A., R.R., J.E., and G.G. interpreted the data.

