## Supplementary Figures for "Novel mechanism of *MYC* deregulation in Multiple Myeloma"

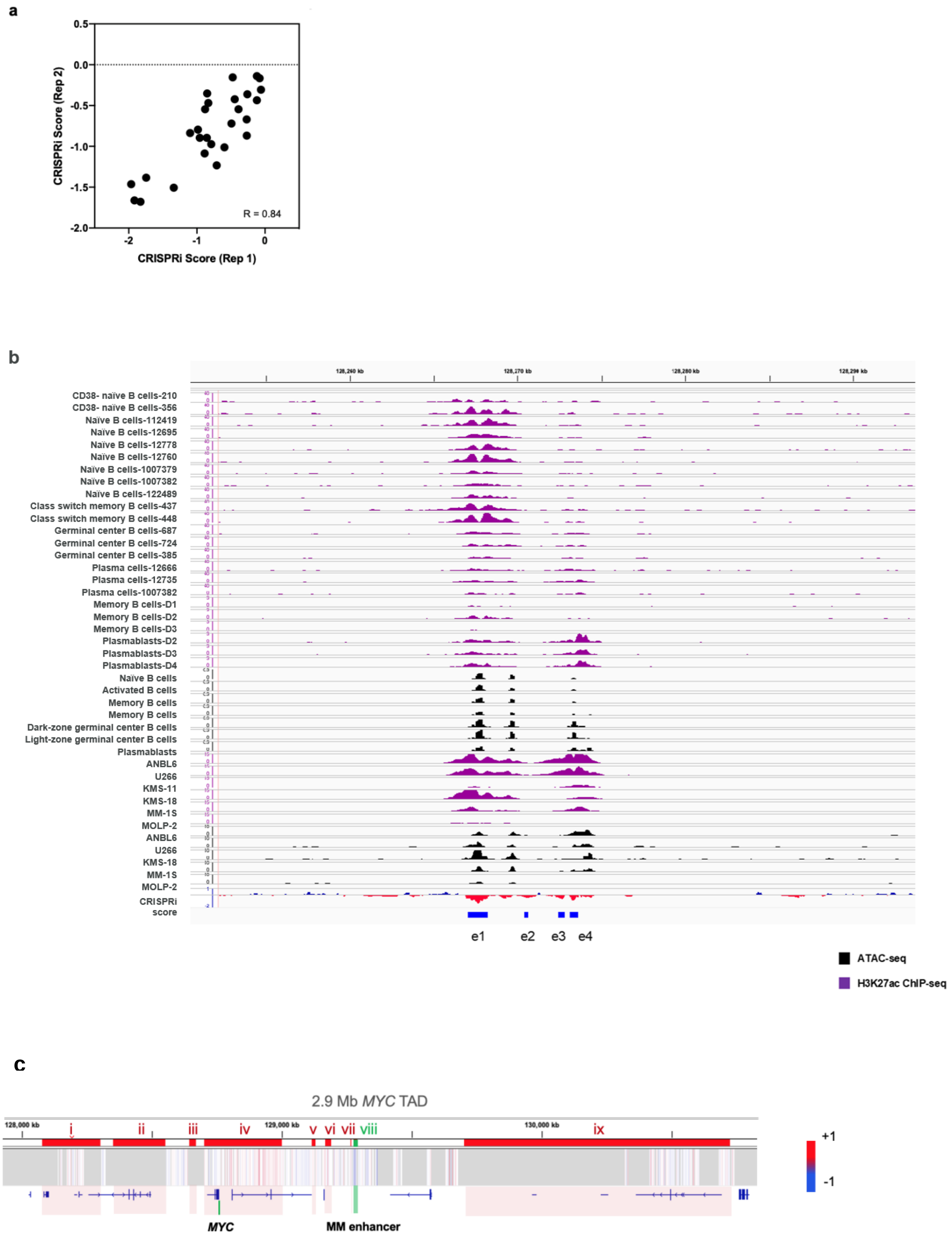
a
CRISPRi Score (Rep 2)
CRISPRi Score (Rep 1)
R = 0.84
0.5
0.0
-0.5
-1.0
-1.5
-2.0
-2
-1
0
b
128,260 kb
128,270 kb
128,280 kb
128,290 kb
CD38- naïve B cells-210
CD38- naïve B cells-356
Naïve B cells-112419
Naïve B cells-12695
Naïve B cells-12778
Naïve B cells-12760
Naïve B cells-1007379
Naïve B cells-1007382
Naïve B cells-122489
Class switch memory B cells-437
Class switch memory B cells-448
Germinal center B cells-687
Germinal center B cells-724
Germinal center B cells-385
Plasma cells-12666
Plasma cells-12735
Plasma cells-1007382
Memory B cells-D1
Memory B cells-D2
Memory B cells-D3
Plasmablasts-D2
Plasmablasts-D3
Plasmablasts-D4
Naïve B cells
Activated B cells
Memory B cells
Memory B cells
Dark-zone germinal center B cells
Light-zone germinal center B cells
Plasmablasts
ANBL6
U266
KMS-11
KMS-18
MM-1S
MOLP-2
ANBL6
U266
KMS-18
MM-1S
MOLP-2
CRISPRi score
e1
e2
e3
e4
ATAC-seq
H3K27ac ChIP-seq
c
2.9 Mb MYC TAD
128,000 kb
129,000 kb
130,000 kb
i
ii
iii
iv
v
vi
vii
viii
ix
+1
-1
MYC
MM enhancer

**Supplementary Fig. 1. Newly identified enhancer region is MM-specific. a)** CRISPRi score for 27 sgRNAs targeting *MYC* TSS in replicate screen, **b)** Chromatin accessibility in a 48 kb sequence around *MYC* enhancer region (hg38) in naïve and activated B cells, germinal center and memory B cells, plasmablasts and plasma cells and MM cell lines, **c)** Locations of known enhancer clusters previously shown to regulate *MYC* expression in other cancers: i) prostate cancer, B cell malignancies (CLL, MCL), ii) breast cancer, prostate cancer, colorectal cancer, iii) medulloblastoma, iv) MCC, v) neuroblastoma, vi) lung cancer, vii) AML, viii) newly identified MM enhancer region, ix) uterine and ovarian cancer, T-ALL, Neuroblastoma, Glioma, AML. The heatmap of CRISPRi scores in the screen of MM cells displayed only at the novel enhancer sequence.

**a**

**b**

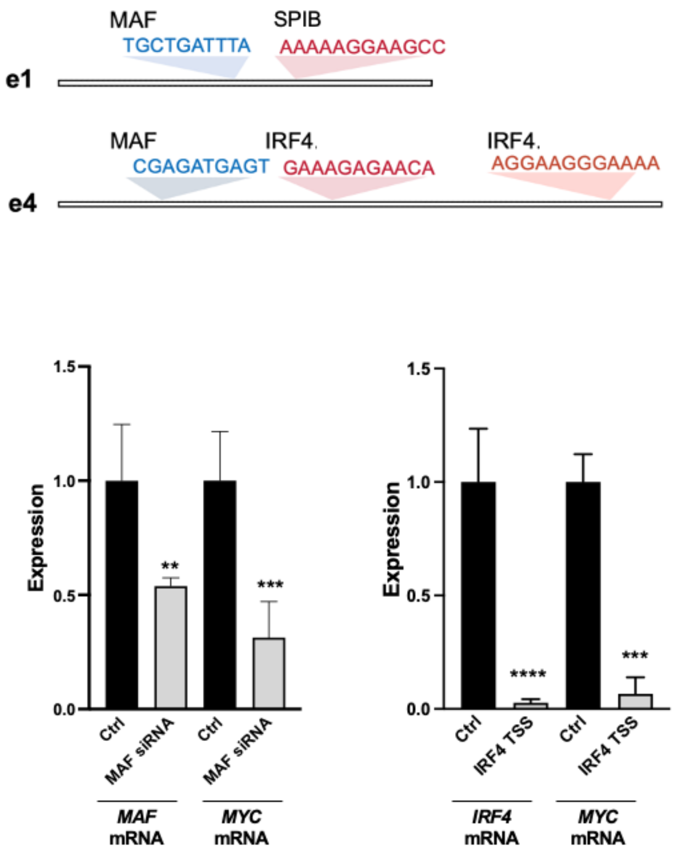


**Supplementary Fig. 2. IRF4 and cMAF are required for *MYC* expression. a**) IRF4, SPIB and cMAF motifs discovered on e1 and e4 enhancers. **b)** siRNA and sgRNA knock down of cMAF and IRF4 in ANBL6 cells. Cells were harvested 48 hours after transfection and *MYC* mRNA levels were measured by real-time PCR. *MYC* expression values were normalized on control samples. Error bars represent 95% CI for the mean values of *MYC* expression values in each sample. Black bars: *MYC* expression levels in control cells (Ctrl) transfected with siRNA and sgRNAs targeting no region in the genome. Gray bars: *MYC* expression in IRF4 or cMAF knocked down cells. ** $P < 0.01$, *** $P < 0.001$, **** $P < 0.0001$ in t-test versus Ctrl sgRNAs or siRNAs.

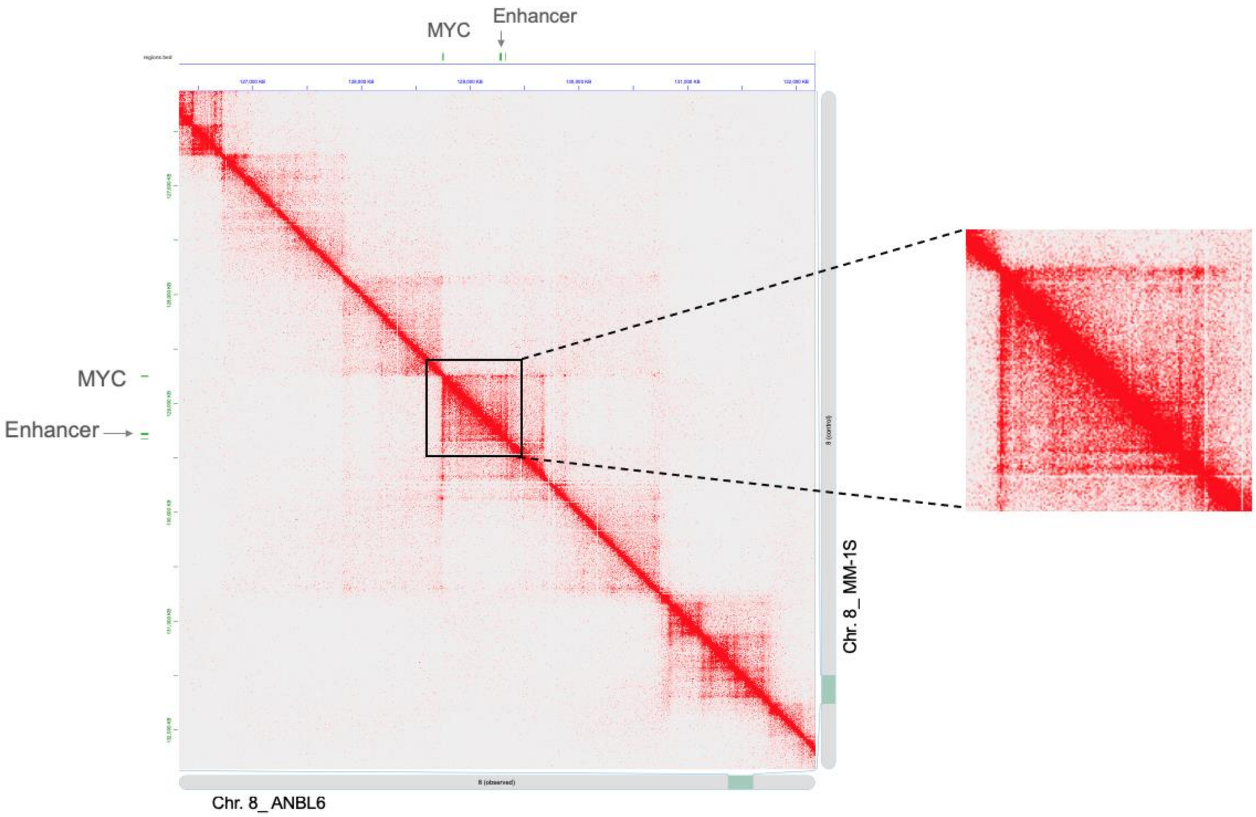


**Supplementary Fig. 3. *MYC* enhancer-promoter interactions in MM cells with active enhancer.** Hi-C contact map showing pairwise genomic contact frequencies. Contacts are shown between genomic coordinates are shown on the x and y axes as a heatmap where red color indicates pairwise contacts. The genomic locations of the *MYC* promoter and enhancer are shown, and higher intra domain *MYC* promoter-enhancer interactions in ANBL6 compared to MM1s is highlighted in the magnified view to the right.
