## Supplementary Tables for "Novel mechanism of *MYC* deregulation in Multiple Myeloma"

**Supplementary Table 1.** Clinical characteristics of patients whose samples were analyzed by ATAC-sequencing

| Patient ID | Age/Sex | Ig usage | Chromosomal abnormalities |
|---|---|---|---|
| MM1 | 68/ Female | IgG/ Kappa | HRD |
| MM2 | 72/ Male | IgG/ Kappa | HRD |
| MM3 | 65/ Male | IgG/ Lambda | t(4;14), 1q duplication, Monosomy 13 |
| MM4 | 54/ Male | IgG/ Kappa | HRD, MYC translocation |
| MM5 | 70/ Male | IgG/ Kappa | 1q amplification, t(11;14), MYC translocation |

HRD = Hyperdiploid

**Supplementary Table 2.** Sequences of sgRNAs and siRNAs and qPCR primers used in this study.

| Experiment | sgRNA target | sgRNA | sgRNA Sequence |
|---|---|---|---|
| CRISPRi | Negative Control | 1 | GATCGCGAGGACCCGTTCCGCC |
| CRISPRi | Negative Control | 2 | GACTCGTCACATGGGGTTGCGA |
| CRISPRi | Negative Control | 3 | GACGGAGGAAGTACACAGCT |
| CRISPRi | Negative Control | 4 | GGAGAGGCCCCTGTCGCGT |
| CRISPRi | Negative Control | 5 | GATTGGTTAGGAGAGTGTGTAT |
| CRISPRi | MYC-TSS | 1 | GCTGTAGTAATTCCAGCGAG |
| CRISPRi | MYC-TSS | 2 | GCGCTGCGGGCGTCCTGGGAA |
| CRISPRi | MYC-e1 | 1 | GACCCACACAGAGTGACCATG |
| CRISPRi | MYC-e1 | 2 | GAGCTCTAGAGCGGCATGCAA |
| CRISPRi | MYC-e2 | 1 | GCCATGCCTGCTCCTTAAAGA |
| CRISPRi | MYC-e2 | 2 | GACAGGATGGCAGAATTTGCT |
| CRISPRi | MYC-e3 | 1 | GCAGATACCATCGCAGAGCTC |
| CRISPRi | MYC-e3 | 2 | GCTGGTAATGCAAGTGCTCTT |
| CRISPRi | MYC-e4 | 1 | GATGCCAGGACCTCTCACCTC |
| CRISPRi | MYC-e4 | 2 | GAGGTCTCTAAGTGTAGTCGC |
| CRISPRi | IRF4 | 1 | GGTACTTGCCGCTGTCGATC |
| CRISPRi | IRF4 | 2 | CAAGCAGGACTACAACCGCG |
| CRISPR KO | e1-SPIB | 1 | AGGAGGAAAAAAAGGAAGCCTG |
| CRISPR KO | e1-SPIB | 2 | AGAATGAAGGAGGAAAAAAAG |
| CRISPR KO | e4-IRF4 | 1 | ACATTCAAATTAAAAAGGAAGG |
| CRISPR KO | e4-IRF4 | 2 | ATGCGGTTTTGTTTCTCTTTCG |
| CRISPR KO | e4-cMAF | 1 | TGAATTTTCGAGATGAGTCAG |
| CRISPR KO | e4-cMAF | 2 | TTTGAGATGAGTCAGGAATTG |
| siRNA | cMAF-SMARTpool | | siGENOME-Horizon discovery |
| siRNA | Non-targeting pool #1 | | siGENOME-Horizon discovery |

| Locus | Forward Primer | Reverse Primer |
|---|---|---|
| MYC | TCCCTCCACTCGGAAGGAC | CTGGTGCATTTTCGGTTGTTG |
| CCND2 | CTTCCGCAGTGCTCCTACTT | CCAAGAAACGGTCCAGGTAA |
| CDC25A | GTCGTGAAGGCGCTATTTG | TGAATCTGTTGACTCGGAGGA |
| RIOX2 | TCATGTCGGGCCTAAGAGAC | GGCATTTGATTCTGCAAAGG |
| GAPDH | AGCACATCGCTCAGACAC | GCCCAATACGACCAAATCC |
| c-MAF | GCTTCCGAGAAAACGGCTC | TGCGAGTGGGCTCAGTTATG |
